# Opioid Antagonism in Humans: A Primer on Optimal Dose and Timing for Central Mu-Opioid Receptor Blockade

**DOI:** 10.1101/2022.02.25.481943

**Authors:** Martin Trøstheim, Marie Eikemo, Jan Haaker, J. James Frost, Siri Leknes

## Abstract

Non-human animal studies outline precise mechanisms of central mu-opioid regulation of pain, stress, affiliation and reward processing. In humans, pharmacological blockade with non-selective opioid antagonists such as naloxone and naltrexone is typically used to assess involvement of the mu-opioid system in such processing. However, robust estimates of the opioid receptor blockade achieved by opioid antagonists are missing. Dose and timing schedules are highly variable and often based on single studies. Here, we provide a detailed analysis of central opioid receptor blockade after opioid antagonism based on existing positron emission tomography data. We also create models for estimating opioid receptor blockade with intravenous naloxone and oral naltrexone. We find that common doses of intravenous naloxone (0.10-0.15 mg/kg) and oral naltrexone (50 mg) are more than sufficient to produce full blockade of central MOR (>90% receptor occupancy) for the duration of a typical experimental session (∼60 minutes), presumably due to initial super saturation of receptors. Simulations indicate that these doses also produce high KOR blockade (78-100%) and some DOR blockade (10% with naltrexone and 48-74% with naloxone). Lower doses (e.g., 0.01 mg/kg intravenous naloxone) are estimated to produce less DOR and KOR blockade while still achieving a high level of MOR blockade for ∼30 minutes. The models and simulations form the basis of two novel web applications for detailed planning and evaluation of experiments with opioid antagonists. These tools and recommendations enable selection of appropriate antagonists, doses and assessment time points, and determination of the achieved receptor blockade in previous studies.

## 1. Introduction

A variety of psychological processes are thought to be modulated by the brain’s mu-opioid system, including reward [1], pain [2], stress [3], and social bonding [4,5]. A popular method to study the mu-opioid system in humans is the pharmacological blockade of opioid receptors with antagonistic drugs. Opioid receptor antagonists bind opioid receptors, but in contrast to agonists they do not generally produce a response by the cell (although some, e.g., naloxone, may act as inverse agonists under certain conditions [6]). Opioid antagonists such as naloxone and naltrexone have a high affinity to mu-opioid receptors (MOR) and thereby prevent other ligands (including endogenous ones) from binding to this receptor type. Therefore, when antagonism with these drugs blocks a behavior, the behavior is assumed to be mu-opioid-dependent [7]. Most opioid antagonists available for human use are non-selective for opioid receptor subtypes and also bind to kappa-opioid receptors (KOR) with high affinity and to delta-opioid receptors (DOR) with low affinity (Table S1). To enable causal inferences about mu-opioid receptor functions based on pharmacological blockade, it is optimal to select an antagonist, a dose, and an assessment time point that results in complete blockade of MOR while causing minimal interference with other receptor types.

Antagonist doses used in basic human research to block the mu-opioid system are often based on plasma concentration, estimates from single positron emission tomography (PET) studies, or on conventions (e.g., 0.10-0.15 mg/kg intravenous naloxone and 50 mg oral naltrexone), and they vary considerably. For example, reported intravenous naloxone doses used in studies of endogenous mu-opioid function are as low as 0.006 mg/kg [8] and as high as 6 mg/kg [9]. Concurrent KOR and DOR blockade is seldom considered when selecting doses.

PET and dual-detector systems use radiolabeled ligands to quantify in vivo receptor binding in the human brain [10]. Because antagonist drugs prevent accumulation of the radiotracer in the brain, positron emission-based techniques can also be used to estimate receptor blockade with these drugs (Supplement) [11,12]. Here we synthesize the available PET and dual-detector data and create models for estimating the amount and duration of central opioid receptor blockade with various doses of commonly used opioid antagonists. In line with previous interpretations of blockade estimates, we define full blockade as >90% receptor occupancy [13,14]. This overview will help determine the achieved MOR blockade in previous studies and evaluate the possibility of DOR and KOR blockade or carry-over effects affecting the results or complicating inferences. It will also enable selection of the appropriate antagonist drugs, doses, assessment time points and intersession intervals for future studies.

## 2. Materials and Methods

We synthesized and further analyzed the available evidence from PET and dual-detector studies on opioid receptor blockade with naloxone and naltrexone. This project used data extracted from published articles and did therefore not require ethical approval. Studies containing blockade data were located using a semi-systematic approach, based on Web of Science searches and examination of references in relevant papers (Supplement). Model specification, fitting, and diagnostics were performed in R [15] using the packages minpack.lm [16], investr [17], nlstools [18], miceNls [19], qpcR [20], and aomisc [21]. First, we used non-linear least squares regression to model the relationship between antagonist dose and MOR blockade. Using data on MOR blockade half-life, we adjusted blockade estimates from the dose-blockade model according to a specified time since administration of an opioid antagonist. Next, we used the linpk package [22] and data on MOR blockade half-life and time-to-peak MOR blockade to describe the time-blockade relationship. Finally, we simulated DOR and KOR blockade from MOR blockade using the antagonists’ average receptor affinities (Table S1).

The available data on central opioid receptor blockade with nalmefene and GSK1521498 are summarized in the Supplement (Table S1, Figure S2-S3).

### 2.1. Modeling the dose-blockade relationship

Following Mayberg and Frost [13] and Rabiner et al. [23], we fitted a log-logistic model to the available data to describe the relationship between antagonist dose and central blockade at the measurement time point (*t*_*measure*_). We specifically used a four-parameter model (Equation 1) with the lower limit constrained to 0, the upper limit constrained to 100, and the Hill slope constrained to 1. The parameter *ED*_*50*_ (i.e., effective dose 50) in this model indicates the estimated dose at which 50% of the receptors are blocked [23].

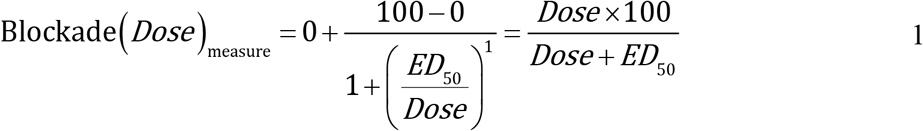

A correction factor was applied to the dose-blockade curve to account for the delay between antagonist administration and MOR blockade assessment in the included studies (Equation 2). This enabled extrapolation of MOR blockade at the administration time point (*t*_*admin*_) assuming no absorption phase. Because MOR blockade with naloxone and naltrexone is eliminated exponentially [24,25], we used an exponential decay model for the correction (Supplement). The elimination rate constant *k* in this model was calculated from available estimates of MOR blockade half-life (Supplement, Equation S3-S7), and the time *t* was set to 0 −*t* _*measure*_ to reflect the temporal delay.

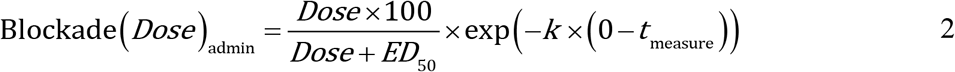

### 2.2. Describing the time-blockade relationship

To describe blockade over time, we adapted the pkprofile function from the linpk package. The pkprofile function is a general pharmacokinetic model for calculating the concentration-time profile of a drug that can account for absorption, infusion duration, and administration of multiple doses [22]. It uses a *V* and *Cl* parameterization where *V* is the volume of distribution and *Cl* is the clearance. *Cl* can be expressed as *k* ×*V*, with *k* being the elimination rate constant calculated from the blockade half-life (see above). By setting *V* to 1, *Cl* defaults to *k* and we can substitute the dose input in the pkprofile function with the estimated MOR blockade at *t*_*admin*_ (Equation 2), assuming no absorption phase. Absorption can instead be handled by the pkprofile function by specifying an absorption rate constant (*k*_*a*_) which can be calculated from the elimination rate constant *k* and the time-to-peak blockade (*t*_*max*_).

The time-to-peak blockade was estimated from time series data of [^11^C]carfentanil activity in the absence and presence of an antagonist. To obtain the absorption rate constant *k*_*a*_ that results in the MOR blockade-time profile of a bolus dose peaking at *t*_*max*_, we used the Lambert W function (*W*_*-1*_) implemented in the pracma package [26] (Supplement, Equation S10-S17).

The time-blockade profiles produced by the pkprofile function treat blockade as a truncated measure of the concentration of the antagonist in the brain. While blockade has an upper limit of 100%, the central concentration may exceed the level necessary to produce full MOR blockade. We allowed model estimates to exceed 100% to reduce underestimation of the duration of full MOR blockade and facilitate detection of excessive concentration levels that contribute to high DOR and KOR blockade.

### 2.3. Simulating delta- and kappa-opioid receptor blockade

Due to limited availability of PET and dual-detector data, we simulated blockade of other opioid receptor subtypes. DOR and KOR blockade was simulated from the MOR blockade by multiplying *ED*_*50*_ for MOR blockade with the affinity of the antagonist drugs to MOR relative to DOR and KOR (see e.g., [27]). We specifically used the relative average affinity from studies of cloned human opioid receptors expressed on Chinese hamster ovary (CHO) cells (Table S1). Time profiles of DOR and KOR blockade assumed similar absorption and elimination rate as for blockade of MOR. Simulation-based estimates were compared to the available PET and dual-detector data on DOR and KOR blockade with naloxone and naltrexone.

## 3. Results

### 3.1. Naloxone

#### 3.1.1. Mu-opioid receptor blockade with intravenous naloxone

Two PET [12,13] and two dual-detector studies [24,28] have used [^11^C]carfentanil to quantify MOR blockade with intravenous naloxone (Table 1). One PET study used a single dose of intravenous naloxone [12] while the other used four different doses [13]. The dual-detector studies used two [24] and eight [28] different doses. Due to the similarities in the protocols used across these studies, we considered the data suitable for quantitative synthesis (Supplement). Timing information was available from a dual-detector study with intravenous naloxone [24] and a PET study with intranasal naloxone [29].

**Table 1.** Overview of positron emission studies used for modeling the dose-blockade relationship of intravenous naloxone and oral naltrexone.

| Antagonist | Method | N | Measurement time point | Dose | MOR blockade |
| --- | --- | --- | --- | --- | --- |
| ***Intravenous naloxone*** |  |  |  |  |  |
| Frost et al. (1985) [12] | PET | 1 | 35-65 minutes | 1 mg/kg | 83%<br>90% |
| Mayberg and Frost (1990) [13] | PET | --- | 45-65<sup>1</sup> minutes | 0.001 mg/kg<br>0.01 mg/kg<br>0.1 mg/kg<br>1.0 mg/kg | 19%<br>65%<br>97%<br>98% |
| Kim et al. (1997) [24] | Dual-detector | 8 | 45-65 minutes | 0.002 mg/kg<br>0.03 mg/kg | 43%<br>81% |
| Villemagne et al. (1994) [28] | Dual-detector | 24 | 45-65 minutes | 0 mg/kg<br>0.0005 mg/kg<br>0.001 mg/kg<br>0.005 mg/kg<sup>1</sup><br>0.01 mg/kg<br>0.1 mg/kg<br>0.5 mg/kg<sup>1</sup><br>1.0 mg/kg | 0%<br>20%<br>40%<br>75%<br>100%<br>100%<br>100%<br>100% |
| ***Oral naltrexone*** |  |  |  |  |  |
| Rabiner et al. (2011) [23] | PET | 20 | < 8 hours | 2 mg<br>5 mg<br>5 mg<br>5 mg<br>15 mg<br>50 mg<br>50 mg | 27%<br>43%<br>46%<br>58%<br>61%<br>92%<br>98% |

*Note.* --- = not reported. <sup>1</sup> Information provided by J. James Frost.

##### 3.1.1.1. Modeling the dose-blockade relationship

Blockade estimates in available PET and dual-detector studies (Table 1) are based on the mean signal recorded between 45-65 minutes after intravenous naloxone administration. Assuming a linear decrease in blockade within this 20-minute time window (Supplement, Figure S1), the reported blockade estimates approximately correspond to the blockade observed halfway (i.e., 10 minutes) through this section of the recording, i.e., at 55 minutes after intravenous naloxone administration (*t*_*measure*_). Figure 1 displays the relationship between dose and MOR blockade (in black) for intravenous naloxone ∼55 minutes after administration. For the log-logistic model (*RMSE* = 9.44; *Pseudo-R*^*2*^ = 0.92; Shapiro-Wilk test: *W* = 0.96, *p* = 0.62; Levene’s test: *F*_*1, 14*_ = 0.06, *p* = 0.81), we obtained the parameter estimate *ED*_*50*_ (*SE*) = 0.0023 mg/kg (0.0004). Although dual-detector studies tended to report higher blockade estimates than PET studies, the difference in *ED*_*50*_ between the two methods was not statistically significant (Δ*ED*_*50*_ = 0.0032 mg/kg, *SE* = 0.0016, *t*_*14*_ = 2.05, *p* = 0.06; Supplement, Equation S1 and S2).

**Figure 1.**
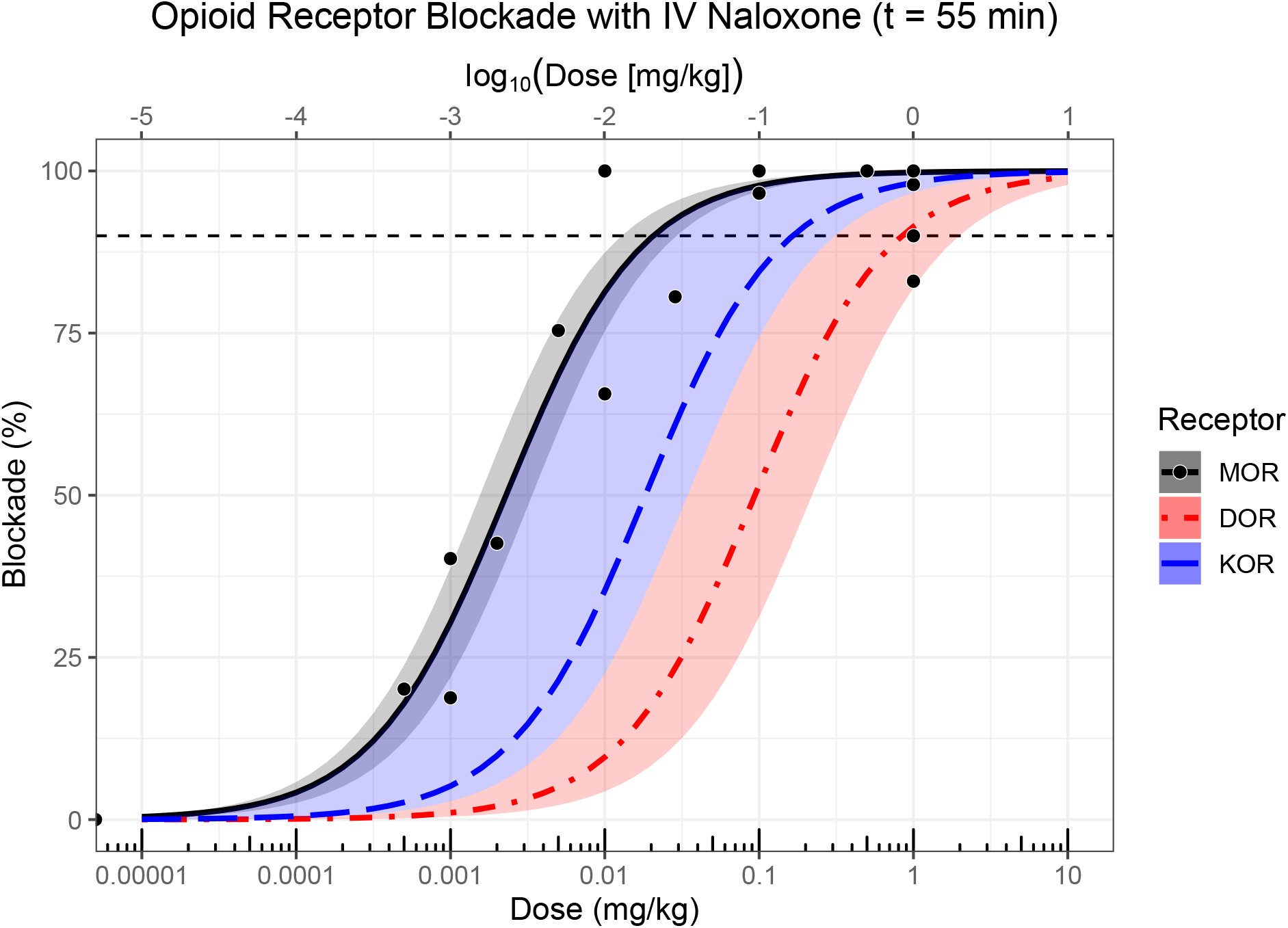
The effect of intravenous (IV) naloxone dose on opioid receptor blockade ∼55 minutes after administration. The bottom x-axis displays untransformed doses while the top x-axis shows the corresponding log_10_-transformed doses. The dashed horizontal line indicates full (90%) receptor blockade. MOR blockade (solid black curve) is based on the data in Table 1 (black dots). DOR (dot-dashed red curve) and KOR blockade (long-dashed blue curve) were approximated from MOR blockade using the relative receptor affinities of naloxone (Table S1). Semitransparent bands indicate 95% confidence band (black band), or range based on highest and lowest reported affinity ratio (blue and red bands; Table S1). The estimated *ED*_*50*_ for MOR, DOR and KOR blockade was 0.0023 (*SE* = 0.0004), 0.094 and 0.018 mg/kg, respectively.

Blockade half-life estimates for naloxone obtained in PET [29] and dual-detector studies [24] are 100 and 120 minutes respectively. Using an average blockade half-life of 110 minutes (*SE* = 10), we obtained the elimination rate constant *k* = 0.006 (Supplement, Equation S18) and the following adjusted formula for calculating MOR blockade as a function of dose at *t*_*admin*_ assuming no absorption phase:

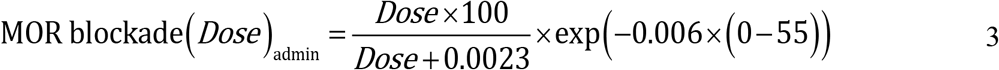

##### 3.1.1.2. Describing the time-blockade relationship

Time series data from dual-detector studies with [^11^C]carfentanil [11,24,25] show a maximum reduction in activity from the control condition (i.e., no naloxone) at 23-29 minutes after administration of intravenous naloxone. When naloxone was administered 5 minutes before [^11^C]carfentanil, maximum signal reduction occurred 29 minutes later with 2 mg naloxone [24], and 23 [11,25], and 26 [24] minutes later with 1 mg/kg naloxone (*M* = 24.7, *SE* = 1.2; Supplement). This suggests that it takes ∼25 minutes after *t*_*admin*_ for naloxone to be distributed to the brain and occupy a maximum amount of central MOR after a single intravenous bolus of naloxone. With the elimination rate constant *k* = 0.006, and *t*_*max*_ = 25, we get an absorption rate constant of *k*_*a*_ = 0.126 (Equation S19). Together with the adjusted dose-dependent blockade estimate at *t*_*admin*_, these timing-based parameters enable estimation of time-blockade profiles of various bolus doses of intravenous naloxone with the pkprofile function (Figure 2).

**Figure 2.**
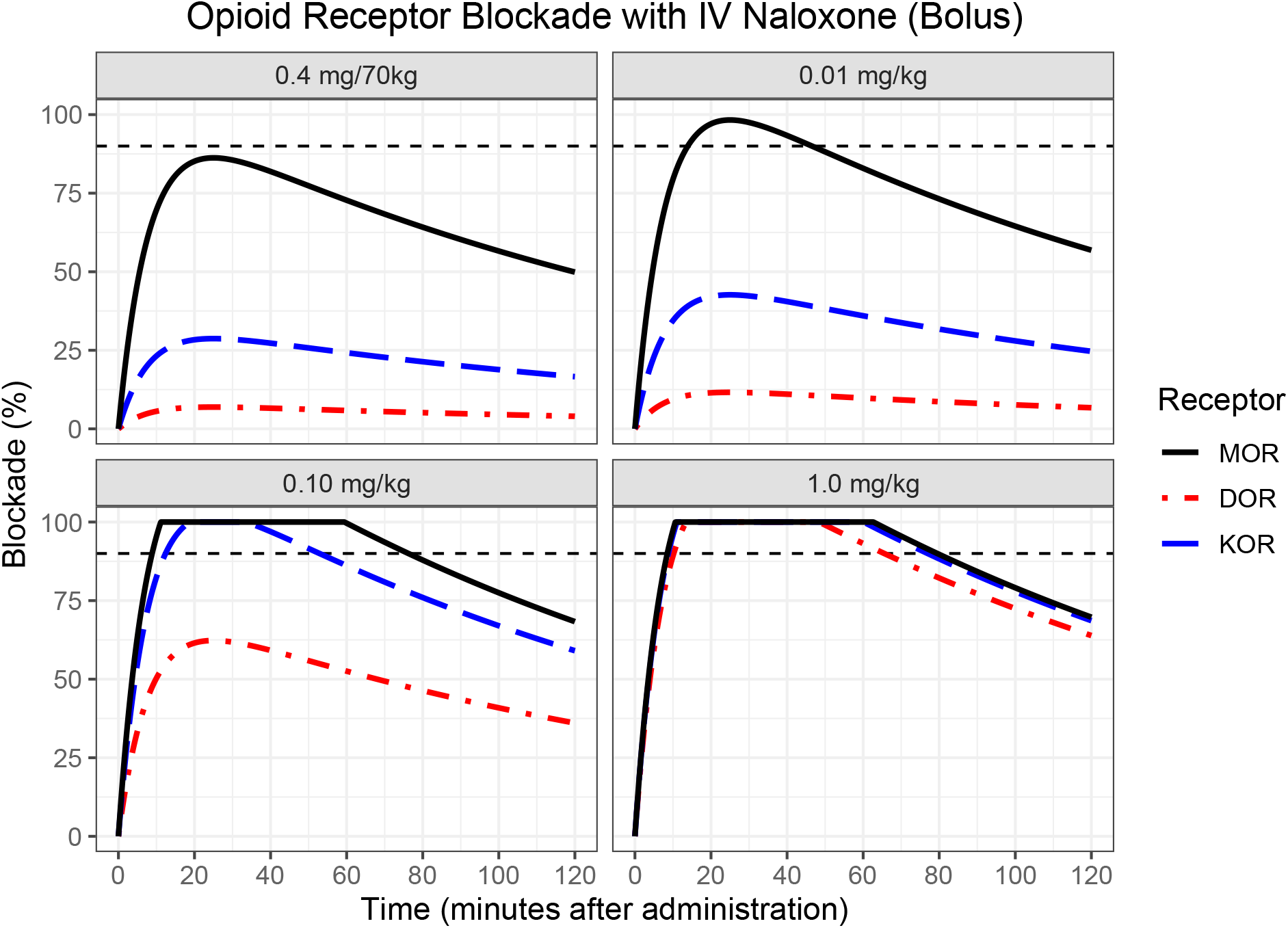
Time-blockade curves for exemplified bolus doses of intravenous (IV) naloxone, accounting for distribution to the brain and truncated at 100% blockade. The dashed horizontal line indicates full (90%) receptor blockade. DOR (dot-dashed red curve) and KOR blockade (long-dashed blue curve) were approximated from MOR blockade (solid black curve) using the relative receptor affinities of naloxone (Table S1).

#### 3.1.2. Delta- and kappa-opioid receptor blockade with intravenous naloxone

Using relative average affinity values (Table S1), we estimated that *ED*_*50*_ would be 41 times greater for DOR than MOR and 8 times greater for KOR than MOR. By multiplying *ED*_*50*_ for MOR blockade with naloxone’s affinity to MOR relative to DOR and KOR, we obtain *ED*_*50*_ = 0.094 mg/kg for DOR blockade and *ED*_*50*_ = 0.018 mg/kg for KOR blockade. This results in the following equations for approximating DOR (Equation 4) and KOR blockade (Equation 5) at *t*_*admin*_, assuming no absorption phase and *k* = 0.006:

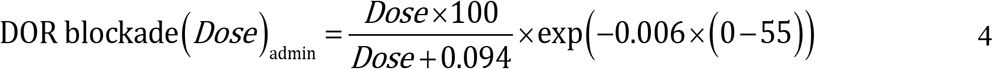

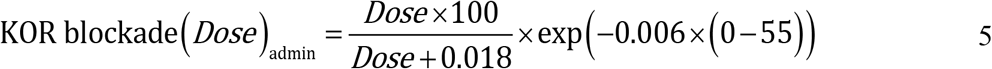

Assuming *k*_*a*_ = 0.126, time-blockade profiles with absorption and eliminations phases can then be simulated with the pkprofile function (Figure 2).

To validate these models, we compared the simulation results to data on DOR and KOR blockade with naloxone in humans. The available data are limited to studies using the non-selective opioid agonist [^11^C]diprenorphine which has equal affinity to MOR, DOR and KOR. These studies suggest that doses of 0.1-1.5 mg/kg intravenous naloxone can completely block all three major opioid receptors [28,30]. A lower dose of ∼0.01 mg/kg produced full MOR blockade, but only partial blockade of DOR/KOR [28,31]. KOR blockade would likely be greater compared to DOR blockade due to naloxone’s higher affinity to KOR (Table S1). Our simulations are largely consistent with the available data, showing full KOR and high DOR blockade with 0.10 mg/kg and partial DOR and KOR blockade with 0.01 mg/kg (Figure 2). However, PET studies with ligands selective to DOR and KOR are necessary to determine the differential blockade of these two receptors by intravenous naloxone.

### 3.2. Naltrexone

#### 3.2.1. Mu-opioid receptor blockade with oral naltrexone

##### 3.2.1.1. Single dose

Several studies have used PET and dual-detection systems to investigate MOR blockade with single doses of oral naltrexone. Approximately 2 hours after administration of 50 mg oral naltrexone, the [^11^C]carfentanil signal in the brain matched the signal recorded 35-65 minutes after intravenous administration of 1 mg/kg naloxone [25], suggesting almost complete blockade of mu-opioid receptors [13]. Consistent with this, Rabiner et al. [23] report that 50 mg oral naltrexone produced 95% mu-opioid receptor blockade within 8 hours after administration. The same dose maintained >90% blockade at ∼49 hours after administration in the study by Lee et al. [25]. The observed blockade in this study decreased to 80% at ∼73 hours, 46% at ∼121 hours, and 30% at ∼169 hours after administration of naltrexone. Based on these data, Lee et al. [25] estimated the blockade half-life of naltrexone in the brain to be 72 hours. Lower doses of oral naltrexone also produce substantial levels of blockade. Within 8 hours of administration, 2, 5 and 15 mg blocked 27, 49 and 61% of the receptors, respectively [23]. Bednarczyk et al. [32] administered 12.5, 50 and 100 mg oral naltrexone and measured blockade after 3, 24, 72 and 144 hours (see also [33]). The blockade estimates from this study are unfortunately unavailable. A study of obese subjects using the non-selective opioid [^11^C]diprenorphine found that a single dose of 150 mg oral naltrexone produced 90% opioid receptor blockade 2 hours after administration [34].

A dose-blockade curve (*RMSE* = 6.96; *Pseudo-R*^*2*^ = 0.92; Shapiro-Wilk test: *W* = 0.98, *p* = 0.93; Levene’s test: *F*_*1, 5*_ = 0.08, *p* = 0.79) based on data obtained within 8 hours of administration was available from Rabiner et al. [23] (Table 1 and Figure 3). This yielded an *ED*_*50*_ of 5.59 mg (*SE* = 0.80) for MOR blockade with oral naltrexone and the following formula for converting dose to blockade:

**Figure 3.**
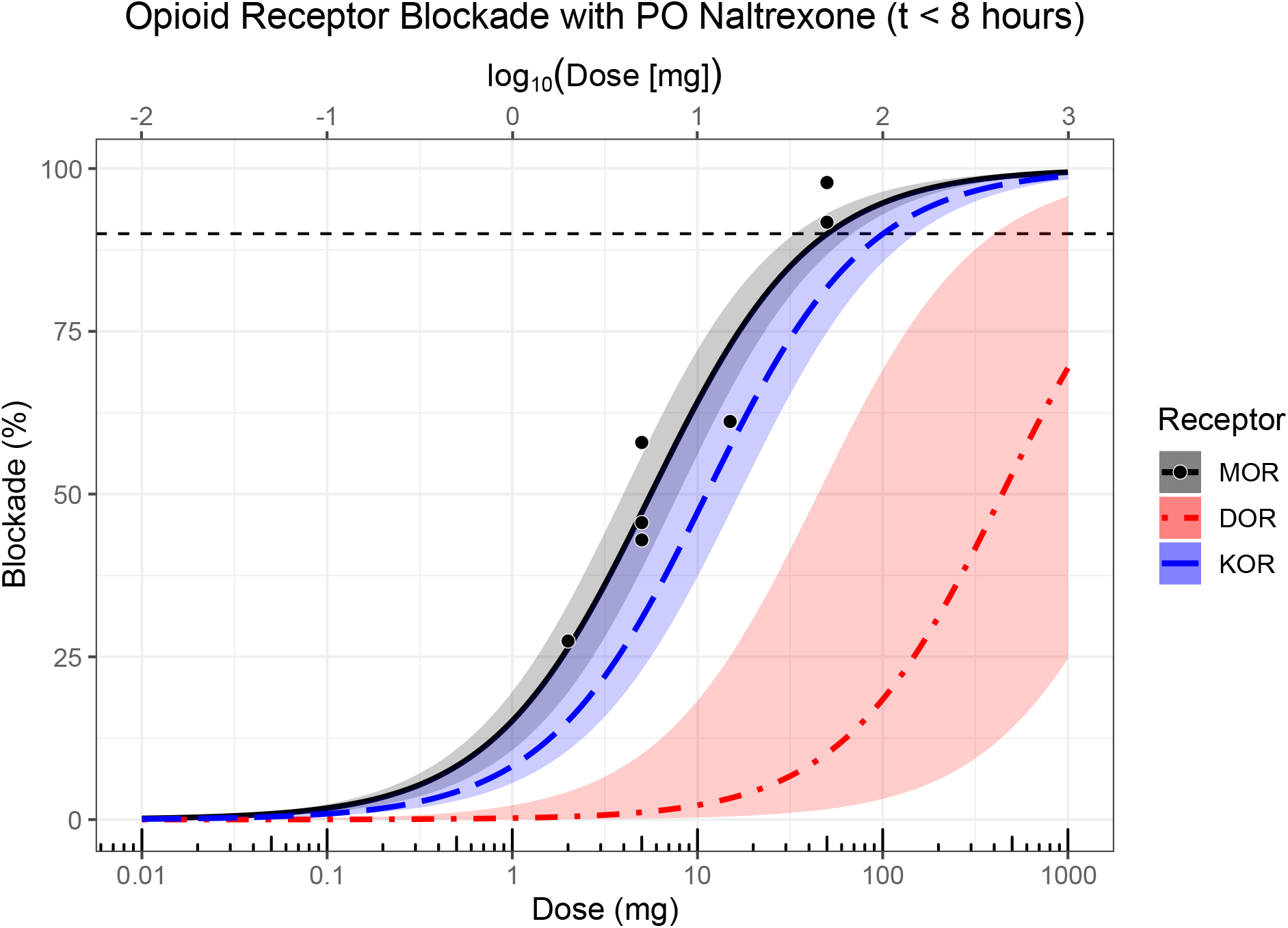
The effect of oral (PO) naltrexone dose on opioid receptor blockade within 8 hours of administration. The bottom x-axis displays untransformed doses while the top x-axis shows the corresponding log_10_-transformed doses. The dashed horizontal line indicates full (90%) receptor blockade. MOR blockade (solid black curve) is based on data (black dots) from Rabiner et al. [23] (Table 1). DOR (dot-dashed red curve) and KOR blockade (long-dashed blue curve) were approximated from MOR blockade using the relative receptor affinities of naltrexone (Table S1). Semitransparent bands indicate 95% confidence band (black band), or range based on highest and lowest reported affinity ratio (blue and red bands; Table S1). The estimated *ED*_*50*_ for MOR, DOR and KOR blockade was 5.59 (*SE* = 0.80; see also [23], 441.83 and 11.19 mg, respectively.

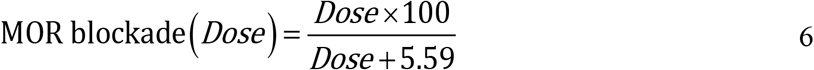

Insufficiently detailed timing information prevented generation of time-blockade profiles for this antagonist.

##### 3.2.1.2. Repeated administration

The effect of repeated naltrexone administration on mu-opioid receptor availability has been investigated with PET in abstinent alcohol dependent patients and in obese subjects. Following four days of treatment with 50 mg oral naltrexone, MOR blockade reached 95% in a sample of abstinent alcohol dependent patients [14]. High levels of MOR blockade (75-97%) was observed in a similar patient sample after daily administration of 50 mg oral naltrexone for three days [35]. In an [^11^C]diprenorphine study of obese subjects who had completed seven days of treatment with oral naltrexone, opioid receptor blockade was 70-80% in those who had received 16 mg/day and 90% in those who had received 32 and 48 mg/day [34]. Due to the non-selectiveness of the radiotracer and naltrexone’s preference for MOR (Table S1), it is possible that the estimated blockade in this study is an underestimations of MOR-specific blockade.

#### 3.2.2. Delta- and kappa-opioid receptor blockade with oral naltrexone

The estimated affinity of naltrexone to MOR was 79 times greater than to DOR and 2 times greater than to KOR (Table S1). Thus, we obtained *ED*_*50*_ = 441.83 mg for DOR blockade and *ED*_*50*_ = 11.19 mg for KOR blockade. Blockade of DOR and KOR within 8 hours of oral naltrexone administration can then be simulated with the following models:

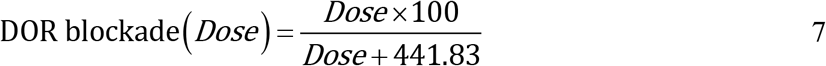

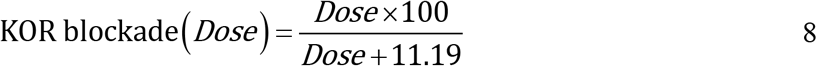

Simulation-based estimates largely agree with results from existing PET studies. For example, studies using the selective KOR agonist [^11^C]GR103545, the preferential KOR agonist [^11^C]EKAP and the preferential KOR antagonist [^11^C]LY2795050 report high KOR blockade (85-93%) in healthy participants and participants with cocaine dependence 2-3 hours after administration of 150 mg oral naltrexone [36–39]. High levels of KOR blockade (87-92%) have also been observed after a week of daily treatment with 100 mg oral naltrexone in participants with alcohol dependence [27,40]. According to simulations based on data from Rabiner et al. [23] and affinity values from studies on cloned human opioid receptors (Table S1), 50 mg oral naltrexone would block 82% of KOR (Figure 3).

Using the selective DOR antagonist N1’-([^11^C]methyl)naltrindole ([^11^C]MeNTI), Madar et al. [41] and Smith et al. [42] reported that a single dose of 50 and 100 mg oral naltrexone produced 38% and 40-95% DOR blockade (respectively) approximately 2 hours after administration in healthy volunteers. Following three and four days of treatment with 50 mg oral naltrexone, the DOR blockade in abstinent alcohol dependent patients was estimated to 31% [35] and 21% [14], respectively. Simulations indicate that 50 mg oral naltrexone would produce only 10% DOR blockade (Figure 3). This underestimation could result from differences in measurement time points between the Rabiner et al. study and the PET studies of DOR blockade, or from some of the latter studies using repeated administration instead of a single dose.

## 4. Discussion

Pharmacological blockade of a receptor system is a common method for probing the function of that receptor system in the human brain. Positron emission techniques yield data on the achieved level of blockade, but for studies of mu-opioid receptors, existing practices vary widely with regards to doses and assessment timing [43]. Here, we have synthesized the available PET and dual-detector data, and created models and web applications for calculating central opioid receptor blockade with the commonly used opioid antagonists naloxone and naltrexone. General recommendations for selecting optimal antagonist drugs, doses and timings in basic human research based on our models and simulations are outlined in Table 2.

**Table 2.**
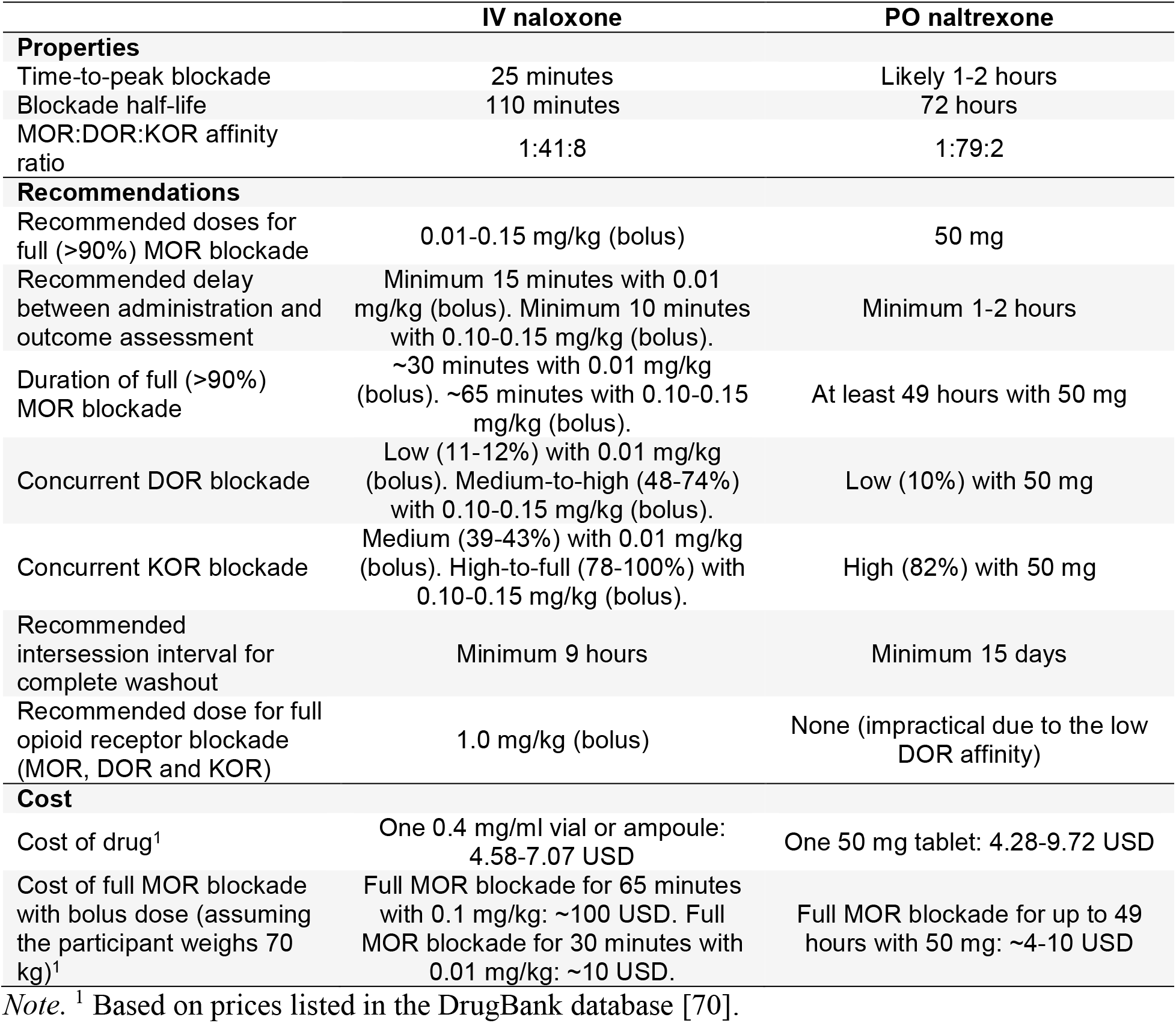
Summary and general recommendations.

To simplify planning of future studies and evaluation of past studies with opioid antagonists, we have designed two web applications using the R package Shiny [44] that incorporate the models for intravenous naloxone (https://martintrostheim.shinyapps.io/planoxone/) and oral naltrexone (https://martintrostheim.shinyapps.io/plantrexone/) presented here. Key features of these applications include estimation of MOR, DOR and KOR blockade over time and across relevant dosing options in human clinical and experimental research, and estimation of total drug amount and cost for planned studies. The source code for both web applications is available on GitHub (https://github.com/martintrostheim/opioid-antagonist-planner).

Pharmacokinetic modeling of opioid antagonists typically focuses on plasma levels. However, for psychopharmacological studies it is important to understand the kinetics of the antagonist in the brain. Available PET and plasma data indicate that the central receptor blockade half-life of intravenous naloxone (110 minutes) and oral naltrexone (72 hours) correspond relatively closely to the plasma half-life of these antagonists during the terminal phase (75 minutes for intravenous naloxone [45] and 96 hours for oral naltrexone [46]), but not during the distribution phase. This suggests that plasma level would be a poor proxy for receptor blockade during the distribution phase and that modeling the elimination from the brain, as approximated here, is needed inform psychopharmacological experiments in sufficient detail.

These novel analytical tools can also aid interpretation of reported effects in the literature, since the presented models yield several insights into how previously used doses affect opioid receptors at the time of assessment. This is especially useful when interpreting the literature using naloxone, which has a relatively short half-life in the brain. Bolus doses of intravenous naloxone used in basic human research are often as large as or larger than 0.10-0.15 mg/kg (e.g., [47–49]). The initial bolus is sometimes supplemented with continuous infusion or an additional bolus (e.g., [49–51]), indicating that many authors may have underestimated the duration of the full blockade with these doses (∼60 minutes). As our model shows, such supplements are only necessary if researchers want to assess the effect of full MOR blockade on outcomes measured more than an hour after the initial bolus (see Figure 3). Yet, testing typically occurs within 15-60 minutes after the initial bolus. Lower doses may be sufficient to produce full MOR blockade, but for a shorter period of time. For example, our model estimates 0.01 mg/kg to maintain full MOR blockade for ∼30 minutes. Combining a low bolus dose with continuous infusion can greatly extend the duration of the full MOR blockade for only a fraction of the cost of a high bolus dose (Table 2). An added benefit of using lower doses is that lower doses typically yield weaker and/or fewer side effects.

Studies using naltrexone to probe the endogenous mu-opioid system often administer 50 mg orally and begin outcome assessment ∼60-120 minutes later (e.g., [52–55]). However, some studies use a higher dose of 100 mg (e.g., [56,57]). The available PET and dual-detector data indicate that 50 mg is more than sufficient to produce full MOR blockade. This dose likely produces central concentration of naltrexone in excess of the dose required to completely block mu-opioid receptors. Weerts et al. [14] observed high level of and low variability in MOR blockade with 50 mg oral naltrexone. However, this ceiling effect could also be a result of the repeated dosing schedule used in this study. Compared to acute doses, repeated dosing would likely cause naltrexone to accumulate, thereby increasing the blockade and extending its duration. This is consistent with the finding that daily administration of an oral naltrexone dose lower than 50 mg (i.e., 32 mg) can result in full opioid receptor blockade [34].

The earliest available MOR blockade estimate with oral naltrexone was collected ∼2 hours after administration and indicates that waiting 2 hours is sufficient to reach full MOR blockade with 50 mg [25]. Considering that naltrexone plasma levels peak 1 hour after oral administration [46], it is possible that the central blockade peaks in less than 2 hours.

Exponential decay processes are considered to be complete after five to ten half-lives [58]. Assuming a blockade half-life of ∼110 minutes with naloxone, we estimate that a washout period of 9 hours should be sufficient to eliminate the MOR blockade. This allows researchers to arrange experimental sessions on consecutive days in within-subjects designs using naloxone. In contrast, naltrexone’s half-life is estimated to 72 hours. After 1 week, which is a common intersession interval for within-subjects designs with oral naltrexone (e.g., [52,53,59]), the MOR blockade with 50 mg is reported to remain at 30%. To ensure complete elimination of the blockade produced by oral naltrexone, a minimum intersession interval of 15 days would be necessary (i.e., five times the 72-hour blockade half-life [58]; see Supplement for further discussion).

Because the opioid antagonists currently marketed for human use are non-selective, there is a concern that DOR and KOR blockade could complicate inferences about MOR functions. The available data and the simulations presented here indicate that naloxone and naltrexone can produce considerable KOR and DOR blockade depending on the dose. Our results need further validation against human PET data as we used highly variable receptor affinity data from CHO cells to simulate human DOR and KOR blockade from MOR blockade. While using a lower naloxone dose could reduce DOR and KOR blockade, this comes at the cost of a shorter duration of full MOR blockade from a bolus injection (see Figure 3). Applying a range of doses optimized for each receptor type can help disentangle the effects of MOR, DOR and KOR blockade on the outcome of interest. For researchers primarily interested in the mu-opioid system, more selective antagonists like GSK1521498 (Supplement) could be a viable alternative to naloxone and naltrexone provided that detailed timing information becomes available.

The models and recommendations presented here should be considered in light of certain limitations. Here, we have used 90% receptor occupancy as the threshold for full blockade [13,14], but we cannot exclude that the up to 10% of unblocked receptors may fulfil some endogenous functions.

While our models, recommendation and web applications are based on data from multiple studies, it is important to note the limited availability of data on opioid receptor blockade with naloxone and naltrexone. Many of the studies tested a small sample of participants, and some blockade estimates are based on a dose applied to a single participant. Due to the variability in design and results between studies, researchers may want to adjust how different data sources are weighted when estimating blockade with our web applications. We have therefore enabled users to tweak all parameters of the models, including *ED*_*50*_, time-to-peak, half-life, and affinity ratios. Data from future PET studies can be used to validate and improve our models and recommendations (see Supplement for further discussion).

The majority of data informing this overview and models were collected from male participants. Men and women are pharmacokinetically and pharmacodynamically different [60], and these differences might affect the opioid receptor blockade produced by antagonist drugs. Many of the studies summarized here either present no analysis of gender effects [13,24,25,28,31,32,35,37,41], tested men only [23,30,36,61], or tested a single participant [11,12]. The few studies that report analyses of gender effects report no significant differences in receptor blockade between men and women [14,27,38,40]. Note that these studies used large doses (e.g., 50-150 mg oral naltrexone) that produce full MOR and KOR blockade in most participants. However, Weerts et al. [14] report no significant relationship between gender and the low DOR blockade produced by daily dosing of 50 mg oral naltrexone. More data is needed determine the generalizability of our models to men and women separately.

The accuracy of opioid receptor blockade estimates with opioid antagonists depends in part on the radiotracer’s affinity to the various opioid receptor subtypes. [^11^C]carfentanil (MOR:DOR:KOR ratio = 1:137:1796)[62], [^11^C]MeNTI (700:1:3250)[63], and [^11^C]GR103545 (810:26800:1)[64] are all highly selective ligands, suggesting that blockade estimates based on the activity of these radiotracers are minimally influenced by their binding to other opioid receptor subtypes. [^11^C]LY2795050 (36:213:1)[65] and [^11^C]EKAP (31:1379:1)[66] preferentially bind to KOR, but they also have affinity to MOR which might influence the KOR blockade estimates obtained with these radiotracer. The non-selectiveness of [^11^C]diprenorphine (1:1:1)[67] makes it difficult to obtain accurate MOR, DOR and KOR blockade estimates with this radiotracer.

Blockade estimates might also be influenced by drug- or disease-related changes to the endogenous opioid system [68]. While most of the reviewed studies included generally healthy volunteers, some included patients with drug dependence [14,27,35,36,40]. Overall, these studies find mixed support for effects of nicotine and alcohol use and dependence on opioid receptor blockade [14,27,40], and no significant relationship between cocaine dependence and opioid receptor blockade [36]. However, the use of large and repeated doses (i.e., 50-150 mg oral naltrexone) in these studies may have resulted in ceiling effects on blockade and thereby prevented the detection of significant predictors. The inconsistent results from an already low number of studies using high and repeated doses highlight the need for more data to determine drug- and disease-related predictors of receptor blockade with opioid antagonists. Such data are key to evaluating the generalizability of our models and recommendations beyond generally healthy people.

The purpose of this primer is to help researchers select antagonist doses that can block endogenous ligands (as modeled by carfentanil) from binding to central opioid receptors. While the information synthesized here may also have implications for the choice of opioid antagonist doses in future clinical research, selection of doses for treatment purposes should primarily be based on knowledge about the relationship between opioid antagonist dose and clinical outcomes. Naloxone and naltrexone are competitive antagonists, meaning that highly potent opioids such as fentanyl, or high doses of opioids like heroin or oxycodone, may overcome the blockade produced by these antagonists. Larger doses or repeated administration of opioid antagonists may therefore be necessary in treatment settings to prevent opioid abuse and to reverse opioid overdose [69].

Pharmacological blockade of the endogenous MOR system with an antagonist such as naloxone and naltrexone is a commonly used method of investigating the role of this system in human psychological processes. While more data on the opioid receptor blockade produced by these antagonists are needed, we hope that this overview and the accompanying tools can aid researchers in evaluating past antagonist studies and in selecting appropriate drugs, doses, assessment time points, and intersession intervals for future studies.

## 5. Funding and Disclosure

This work was supported by grants from the European Research Council under the European Union’s Horizon 2020 research and innovation program (grant agreement No. 802885) to Siri Leknes and the South-Eastern Norway Regional Health Authority (project No. 2018035) to Marie Eikemo. We declare no conflicts of interest.

## 6. Acknowledgements

We thank Howard Fields at the University of California San Francisco, and Molly Carlyle at the University of Oslo and The University of Queensland for feedback on the manuscript. None of the above received financial compensation outside of salary.

## 7. Author contributions

Martin Trøstheim conducted the literature search, extracted and analyzed the data, and wrote the manuscript. J. James Frost contributed with additional data and information. All authors contributed to revising the manuscript.

## Notes

https://github.com/martintrostheim/opioid-antagonist-planner

https://martintrostheim.shinyapps.io/planoxone/

https://martintrostheim.shinyapps.io/plantrexone/

### Competing Interest Statement

The authors have declared no competing interest.

### Summary of Updates

The structure of the manuscript has been updated. The methods and discussion sections have been expanded with additional details and limitations.

## References

1. Meier IM, Eikemo M, Leknes S. The Role of Mu-Opioids for Reward and Threat Processing in Humans: Bridging the Gap from Preclinical to Clinical Opioid Drug Studies. Curr Addict Rep. 2021;8:306–318.

2. Zubieta J-K, Smith YR, Bueller JA, Xu Y, Kilbourn MR, Jewett DM, et al. Regional Mu Opioid Receptor Regulation of Sensory and Affective Dimensions of Pain. Science. 2001;293:311–315.

3. Valentino RJ, Van Bockstaele E. Endogenous opioids: The downside of opposing stress. Stress Resil. 2015;1:23–32.

4. Løseth G, Ellingsen D-M, Leknes S. State-dependent µ-opioid Modulation of Social Motivation – a model. Front Behav Neurosci. 2014;8:430.

5. Machin AJ, Dunbar RIM. The brain opioid theory of social attachment: a review of the evidence. Behaviour. 2011;148:985–1025.

6. Berg KA, Clarke WP. Making Sense of Pharmacology: Inverse Agonism and Functional Selectivity. Int J Neuropsychopharmacol. 2018;21:962–977.

7. Eikemo M, Løseth GE, Leknes S. Do endogenous opioids mediate or fine-tune human pain relief? PAIN. 2021;162:2789–2791.

8. Brennum J, Kaiser F, Dahl JB. Effect of naloxone on primary and secondary hyperalgesia induced by the human burn injury model. Acta Anaesthesiol Scand. 2001;45:954–960.

9. Cutter HSG, O’Farrell TJ. Experience with alcohol and the endogenous opioid system in ethanol analgesia. Addict Behav. 1987;12:331–343.

10. Zhang Y, Fox GB. PET imaging for receptor occupancy: meditations on calculation and simplification. J Biomed Res. 2012;26:69–76.

11. Bice AN, Wagner HN, Frost JJ, Natarajan TK, Lee MC, Wong DF, et al. Simplified Detection System for Neuroreceptor Studies in the Human Brain. J Nucl Med. 1986;27:184–191.

12. Frost JJ, Wagner HNJr, Dannals RF, Ravert HT, Links JM, Wilson AA, et al. Imaging Opiate Receptors in the Human Brain by Positron Tomography. J Comput Assist Tomogr. 1985;9:231–236.

13. Mayberg HS, Frost JJ. Opiate Receptors. In: Frost JJ, Wagner Jr HN, editors. Quant. Imaging Neurorecept. Neurotransmitters Enzym., New York: Raven Press, Ltd.; 1990.

14. Weerts EM, Kim YK, Wand GS, Dannals RF, Lee JS, Frost JJ, et al. Differences in δ- and μ-Opioid Receptor Blockade Measured by Positron Emission Tomography in Naltrexone-Treated Recently Abstinent Alcohol-Dependent Subjects. Neuropsychopharmacology. 2008;33:653–665.

15. R Core Team. R: A language and environment for statistical computing. Vienna, Austria: R Foundation for Statistical Computing; 2020.

16. Elzhov TV, Mullen KM, Spiess A-N, Bolker B. minpack.lm: R Interface to the Levenberg-Marquardt Nonlinear Least-Squares Algorithm Found in MINPACK, Plus Support for Bounds. 2022.

17. Greenwell BM, Kabban CMS. investr: An R Package for Inverse Estimation. R J. 2014;6:90–100.

18. Baty F, Ritz C, Charles S, Brutsche M, Flandrois J-P, Delignette-Muller M-L. A Toolbox for Nonlinear Regression in R: The Package nlstools. J Stat Softw. 2015;66:1–21.

19. Early-Capistrán M-M. miceNls: Utility package for integrating multiple imputation by chained equations (MICE) with nonlinear regression. 2021.

20. Spiess A-N. qpcR: Modelling and Analysis of Real-Time PCR Data. 2018.

21. Onofri A. The broken bridge between biologists and statisticians: a blog and R package. Statforbiology. 2020. https://www.statforbiology.com.

22. Rich B. linpk: Generate Concentration-Time Profiles from Linear PK Systems. 2021.

23. Rabiner EA, Beaver J, Makwana A, Searle G, Long C, Nathan PJ, et al. Pharmacological differentiation of opioid receptor antagonists by molecular and functional imaging of target occupancy and food reward-related brain activation in humans. Mol Psychiatry. 2011;16:826–835.

24. Kim S, Wagner HN, Villemagne VL, Kao P-F, Dannals RF, Ravert HT, et al. Longer Occupancy of Opioid Receptors by Nalmefene Compared to Naloxone as Measured In Vivo by a Dual-Detector System. J Nucl Med. 1997;38:1726–1731.

25. Lee MC, Wagner HN, Tanada S, Frost JJ, Bice AN, Dannals RF. Duration of Occupancy of Opiate Receptors by Naltrexone. J Nucl Med. 1988;29:1207–1211.

26. Borchers HW. pracma: Practical Numerical Math Functions. 2022.

27. de Laat B, Nabulsi N, Huang Y, O’Malley SS, Froehlich JC, Morris ED, et al. Occupancy of the kappa opioid receptor by naltrexone predicts reduction in drinking and craving. Mol Psychiatry. 2021;26:5053–5060.

28. Villemagne VL, Frost JJ, Dannals RF, Lever JR, Tanada S, Natarajan TK, et al. Comparison of [11C]diprenorphine and [11C]carfentanil in vivo binding to opiate receptors in man using a dual detector system. Eur J Pharmacol. 1994;257:195–197.

29. Johansson J, Hirvonen J, Lovró Z, Ekblad L, Kaasinen V, Rajasilta O, et al. Intranasal naloxone rapidly occupies brain mu-opioid receptors in human subjects. Neuropsychopharmacology. 2019;44:1667–1673.

30. Sadzot B, Price JC, Mayberg HS, Douglass KH, Dannals RF, Lever JR, et al. Quantification of Human Opiate Receptor Concentration and Affinity Using High and Low Specific Activity [11C]Diprenorphine and Positron Emission Tomography. J Cereb Blood Flow Metab. 1991;11:204–219.

31. Melichar JK, Nutt DJ, Malizia AL. Naloxone displacement at opioid receptor sites measured in vivo in the human brain. Eur J Pharmacol. 2003;459:217–219.

32. Bednarczyk EM, Wack D, Haka M, Shang Y, Hershey L, O’Sullivan R, et al. Duration of human MU opiate receptor blockade following naltrexone: Measurement by 11C-carfentanil pet. Clin Pharmacol Ther. 2005;77:P26–P26.

33. Okusanya OO, Amer A, Forrest A, Shang E, Bednarczyk EM. Use of PET imaging to develop a pharmacokinetic/pharmacodynamic (PK/PD) model for naltrexone (NTX) & 6-beta-naltrexol (6 beta NTX) occupancy on the human mu-opiate receptor (MOR). Clin Pharmacol Ther. 2007;81:S71.

34. Ye W, Zhou Y, Alexander M, Brasic J, Nandi A, Gruender G, et al. Receptor occupancy following chronic or single daily dosing with naltrexone. J Nucl Med. 2007;48:173P.

35. McCaul ME, Wand GS, Kim YK, Bencherif B, Dannals RF, Frost JJ. Naltrexone Effects on Mu- and Delta-Opioid Receptor Availability in Alcohol Dependence. Alcohol Clin Exp Res. 2003;27:21A.

36. Martinez D, Slifstein M, Matuskey D, Nabulsi N, Zheng M-Q, Lin S, et al. Kappa-opioid receptors, dynorphin, and cocaine addiction: a positron emission tomography study. Neuropsychopharmacology. 2019;44:1720–1727.

37. Naganawa M, Jacobsen LK, Zheng M-Q, Lin S-F, Banerjee A, Byon W, et al. Evaluation of the agonist PET radioligand [11C]GR103545 to image kappa opioid receptor in humans: Kinetic model selection, test–retest reproducibility and receptor occupancy by the antagonist PF-04455242. NeuroImage. 2014;99:69–79.

38. Naganawa M, Zheng M-Q, Nabulsi N, Tomasi G, Henry S, Lin S-F, et al. Kinetic Modeling of 11C-LY2795050, A Novel Antagonist Radiotracer for PET Imaging of the Kappa Opioid Receptor in Humans. J Cereb Blood Flow Metab. 2014;34:1818–1825.

39. Naganawa M, Li S, Nabulsi N, Lin S, Labaree D, Ropchan J, et al. Comparison of 11C-EKAP and 11C-FEKAP, two novel agonist PET radiotracers for imaging the kappa opioid receptor in humans. J Nucl Med. 2017;58:357.

40. Vijay Ai, Morris E, Goldberg A, Petrulli J, Liu H, Huang Y, et al. Naltrexone occupancy at kappa opioid receptors investigated in alcoholics by PET occupancy at kappa opioid receptors investigated in alcoholics by PET. J Nucl Med. 2017;58:1297.

41. Madar I, Lever JR, Kinter CM, Scheffel U, Ravert HT, Musachio JL, et al. Imaging of δ opioid receptors in human brain by N1′-([11C]methyl)naltrindole and PET. Synapse. 1996;24:19–28.

42. Smith JS, Zubieta J-K, Price JC, Flesher JE, Madar I, Lever JR, et al. Quantification of δ-Opioid Receptors in Human Brain with N1′ -([11C]Methyl) Naltrindole and Positron Emission Tomography. J Cereb Blood Flow Metab. 1999;19:956–966.

43. Werner MU, Pereira MP, Andersen LPH, Dahl JB. Endogenous Opioid Antagonism in Physiological Experimental Pain Models: A Systematic Review. PLOS ONE. 2015;10:e0125887.

44. Chang W, Cheng J, Allaire J, Sievert C, Schloerke B, Xie Y, et al. shiny: Web Application Framework for R. 2021.

45. McDonald R, Lorch U, Woodward J, Bosse B, Dooner H, Mundin G, et al. Pharmacokinetics of concentrated naloxone nasal spray for opioid overdose reversal: Phase I healthy volunteer study. Addiction. 2018;113:484–493.

46. Verebey K, Volavka J, Mule SJ, Resnick RB. Naltrexone: Disposition, metabolism, and effects after acute and chronic dosing. Clin Pharmacol Ther. 1976;20:315–328.

47. Bruehl S, Burns JW, Chung OY, Ward P, Johnson B. Anger and pain sensitivity in chronic low back pain patients and pain-free controls: the role of endogenous opioids. PAIN. 2002;99:223–233.

48. Buchel C, Miedl S, Sprenger C. Hedonic processing in humans is mediated by an opioidergic mechanism in a mesocorticolimbic system. ELife. 2018;7:e39648.

49. Eippert F, Bingel U, Schoell ED, Yacubian J, Klinger R, Lorenz J, et al. Activation of the Opioidergic Descending Pain Control System Underlies Placebo Analgesia. Neuron. 2009;63:533–543.

50. Berna C, Leknes S, Ahmad AH, Mhuircheartaigh RN, Goodwin GM, Tracey I. Opioid-Independent and Opioid-Mediated Modes of Pain Modulation. J Neurosci. 2018;38:9047– 9058.

51. Julien N, Marchand S. Endogenous pain inhibitory systems activated by spatial summation are opioid-mediated. Neurosci Lett. 2006;401:256–260.

52. Bruehl S, Carlson CR, Wilson JF, Norton JA, Colclough G, Brady MJ, et al. Psychological coping with acute pain: An examination of the role of endogenous opioid mechanisms. J Behav Med. 1996;19:129–142.

53. Eikemo M, Løseth GE, Johnstone T, Gjerstad J, Willoch F, Leknes S. Sweet taste pleasantness is modulated by morphine and naltrexone. Psychopharmacology (Berl). 2016;233:3711–3723.

54. Inagaki TK, Hazlett LI, Andreescu C. Naltrexone alters responses to social and physical warmth: implications for social bonding. Soc Cogn Affect Neurosci. 2019;14:471–479.

55. Meier IM, Bos PA, Hamilton K, Stein DJ, van Honk J, Malcolm-Smith S. Naltrexone increases negatively-valenced facial responses to happy faces in female participants. Psychoneuroendocrinology. 2016;74:65–68.

56. Charles SJ, Farias M, van Mulukom V, Saraswati A, Dein S, Watts F, et al. Blocking mu-opioid receptors inhibits social bonding in rituals. Biol Lett. 2020;16:20200485.

57. Tarr B, Launay J, Benson C, Dunbar RIM. ‘Naltrexone Blocks Endorphins Released when Dancing in Synchrony’. Adapt Hum Behav Physiol. 2017;3:241–254.

58. Byers JP, Sarver JG. Chapter 10 - Pharmacokinetic Modeling. In: Hacker M, Messer W, Bachmann K, editors. Pharmacology, San Diego: Academic Press; 2009. p. 201–277.

59. Yeomans MR, Gray RW. Effects of Naltrexone on Food Intake and Changes in Subjective Appetite During Eating: Evidence for Opioid Involvement in the Appetizer Effect. Physiol Behav. 1997;62:15–21.

60. Soldin OP, Mattison DR. Sex Differences in Pharmacokinetics and Pharmacodynamics. Clin Pharmacokinet. 2009;48:143–157.

61. Ingman K, Hagelberg N, Aalto S, Någren K, Juhakoski A, Karhuvaara S, et al. Prolonged Central μ-Opioid Receptor Occupancy after Single and Repeated Nalmefene Dosing. Neuropsychopharmacology. 2005;30:2245–2253.

62. Subramanian G, Paterlini MG, Portoghese PS, Ferguson DM. Molecular Docking Reveals a Novel Binding Site Model for Fentanyl at the μ-Opioid Receptor. J Med Chem. 2000;43:381–391.

63. Portoghese PS, Sultana M, Takemori AE. Design of peptidomimetic δ opioid receptor antagonists using the message-address concept. J Med Chem. 1990;33:1714–1720.

64. Schoultz BW, Hjornevik T, Willoch F, Marton J, Noda A, Murakami Y, et al. Evaluation of the kappa-opioid receptor-selective tracer [11C]GR103545 in awake rhesus macaques. Eur J Nucl Med Mol Imaging. 2010;37:1174–1180.

65. Zheng M-Q, Nabulsi N, Kim SJ, Tomasi G, Lin S, Mitch C, et al. Synthesis and Evaluation of 11C-LY2795050 as a κ-Opioid Receptor Antagonist Radiotracer for PET Imaging. J Nucl Med. 2013;54:455–463.

66. Li S, Zheng M-Q, Naganawa M, Kim S, Gao H, Kapinos M, et al. Development and In Vivo Evaluation of a κ-Opioid Receptor Agonist as a PET Radiotracer with Superior Imaging Characteristics. J Nucl Med. 2019;60:1023–1030.

67. Henriksen G, Willoch F, Talbot PS, Wester H-J. Recent development and potential use of µ- and κ-opioid receptor ligands in positron emission tomography studies. Drug Dev Res. 2006;67:890–904.

68. Henriksen G, Willoch F. Imaging of opioid receptors in the central nervous system. Brain. 2008;131:1171–1196.

69. Fairbairn N, Coffin PO, Walley AY. Naloxone for heroin, prescription opioid, and illicitly made fentanyl overdoses: Challenges and innovations responding to a dynamic epidemic. Int J Drug Policy. 2017;46:172–179.

70. Wishart DS, Feunang YD, Guo AC, Lo EJ, Marcu A, Grant JR, et al. DrugBank 5.0: a major update to the DrugBank database for 2018. Nucleic Acids Res. 2018;46:D1074– D1082.

