## Supplement for "Opioid Antagonism in Humans: A Primer on Optimal Dose and Timing for Central Mu-Opioid Receptor Blockade"

### 1. Introduction

PET and dual-detector systems quantify in vivo receptor binding in the human brain and can be used to estimate opioid receptor blockade with non-selective opioid antagonists. Quantification of receptor binding with these techniques works by injecting a radiotracer; a ligand that binds to the target receptors in the brain and that has been labelled with a radioactive isotope. Receptor binding results in accumulation of the radiotracer in the brain. Positron emission from the radioactive isotope can then be recorded through detection of gamma rays produced by electron-positron annihilation [1–3]. Administration of an antagonist prevents the radiotracer from binding to the receptors, resulting in less accumulation of the radiotracer and fewer detected gamma rays.

**Table S1**

*Affinities of the opioid antagonists naloxone, naltrexone, nalmefene, and GSK1521498 to cloned human opioid receptors expressed on Chinese hamster ovary (CHO) cells.*

| | | $K_i$ (nM) | | |
|---|---|---|---|---|
| **Antagonist** | **MOR** | **DOR** | **KOR** | **MOR:DOR:KOR** |
| ***Naloxone*** | | | | |
| Kelly et al. (2015) [4] | 2.09[1] | 30.90[1] | 31.62[1] | 1:15:15 |
| Peng et al. (2007) [5] | 0.79 | 76 | 1.1 | 1:96:1 |
| Toll et al. (1998) [6] | 1.4 | 67.5 | 2.5 | 1:48:2 |
| *Average* (*SE*) | 1.43 (0.38) | 58.13 (13.84) | 11.74 (9.95) | 1:41:8 |
| ***Naltrexone*** | | | | |
| Kelly et al. (2015) [4] | 0.71[1] | 5.50[1] | 1.66[1] | 1:8:2 |
| Konoura et al. (2015) [7] | 0.27 | 12.3 | 0.70 | 1:46:3 |
| Peng et al. (2007) [5] | 0.23 | 38 | 0.25 | 1:165:1 |
| Schüllner et al. (2003) [8] | 0.20 | 8.70 | 0.40 | 1:44:2 |
| Toll et al. (1998) [6] | 0.2 | 10.8 | 0.4 | 1:54:2 |
| Wentland et al. (2005) [9] | 0.11 | 60 | 0.19 | 1:545:2 |
| *Average* (*SE*) | 0.29 (0.09) | 22.55 (8.88) | 0.60 (0.22) | 1:79:2 |
| ***Nalmefene*** | | | | |
| Bart et al. (2005) [10] | 0.24 | 16 | 0.083 | 3:193:1[2] |
| Kelly et al. (20015) [4] | 0.24[1] | 8.32[1] | --- | 1:35:--- |
| Toll et al. (1998) [6] | 0.3 | 7.3 | 0.3 | 1:24:1 |
| *Average* (*SE*) | 0.26 (0.02) | 10.54 (2.75) | 0.19 (0.11) | 1:55:1[2] |
| ***GSK1521498*** | | | | |
| Kelly et al. (2015) [4] | 0.23[1] | 3.16[1] | 3.31[1] | 1:14:14 |
| Rabiner et al. (2011) [11] | 1.5 | 30.2 | 20.4 | 1:20:14 |
| *Average* (*SE*) | 0.87 (0.64) | 16.68 (13.52) | 11.86 (8.55) | 1:19:14 |

*Note.* Lower values indicate higher affinity. --- = not applicable. $K_i$ is the inhibition constant, i.e., the concentration in nanomolar (nM) required to produce half of the maximum blockade of the receptor. As such, lower numbers (i.e., concentrations) thereby indicate higher affinity for the receptor subtype. [1] $K_i$ converted from $pK_i$, i.e., the negative logarithm to base 10 of $K_i$ expressed in molar. [2] Calculated with $K_i$ for KOR as denominator.

### 2. Methods

#### 2.1. Literature search and data extraction

To locate relevant data on opioid receptor blockade with antagonist drugs, we conducted searches in Web of Science using combinations of the terms 1) naloxone, naltrexone, nalmefene or GSK1521498, 2) occupancy, blockade or binding, and 3) positron emission or PET. The searches targeted title, abstract and keywords. We also screened the reference lists of relevant articles to locate additional data. When necessary, data was extracted from figures using WebPlotDigitizer [12]. Digitization with this tool is simple and reliable and results in valid data when compared with the source data [13–15]. We contacted original authors for additional details and missing data.

#### 2.2. Assessing the difference in $ED_{50}$ between PET and dual-detector studies

Dual-detector studies use the [$^{11}$C]carfentanil signal under 1.0 mg/kg intravenous naloxone as a reference for nonspecific binding when calculating MOR blockade. This could result in overestimation of blockade with lower doses in cases when 1.0 mg/kg does produce complete MOR blockade. To assess the influence of recording techniques on MOR blockade estimates, we modified the log-logistic model (Equation S1) so that the $ED_{50}$ term would depend on the recording technique (Equation S2). The recording technique was dummy coded with 0 for dual-detector and 1 for PET.

$$\text{Blockade}(Dose)_{\text{measure}} = \frac{Dose \times 100}{Dose + ED_{50}} \quad \text{S1}$$

$$\text{Blockade}(Dose, Technique)_{\text{measure}} = \frac{Dose \times 100}{Dose + (b_0 + b_1 \times Technique)} \quad \text{S2}$$

In Equation S2, $b_0$ is the intercept, or $ED_{50}$ when the recording technique is the dual-detector system, while $b_1$ is the change in $ED_{50}$ when the recording technique is PET instead of the dual-detector system. We also ran the log-logistic model in Equation S1 separately for PET and dual-detector studies to obtain estimates of $ED_{50}$ for each recording technique.

#### 2.3. Calculating the elimination rate constant from MOR blockade half-life

Because MOR blockade with naloxone and naltrexone is eliminated exponentially [16,17], we can estimate the elimination rate constant from the antagonists' MOR blockade half-life. The below formula (Equation S3) describes the elimination of blockade over time assuming no initial absorption phase.

$$\text{Blockade}(t) = d \times \exp(-k \times t) \qquad \text{S3}$$

The blockade produced by a bolus dose of an antagonist at any given half-life (*h*) can then be expressed as

$$d \times 0.5^{h} = d \times \exp(-k \times t_{1/2} \times h) \qquad \text{S4}$$

To find the elimination rate constant *k* as a function of MOR blockade half-life, we can simply rearrange Equation S4.

$$\ln(0.5^{h}) = \ln(\exp(-k \times t_{1/2} \times h)) \qquad \text{S5}$$

$$h \times \ln(0.5) = -k \times t_{1/2} \times h \qquad \text{S6}$$

$$k = \frac{-\ln(0.5)}{t_{1/2}} = \frac{\ln(2)}{t_{1/2}} \qquad \text{S7}$$

#### 2.4. Extrapolating MOR blockade at the administration time point

We used an exponential decay model with the obtained elimination rate constant *k* (Equation S3) to extrapolate MOR blockade at the administration time point ($t_{admin}$). In Equation S3, *t* is the time in minutes after the measurement time point ($t_{measure}$) and *d* is the dose-dependent blockade at $t_{measure}$, which can be substituted with Equation S1. By subtracting $t_{measure}$ from the time input $t$, we enable extrapolation of blockade between $t_{admin}$ (i.e., 0 minutes after administration) and $t_{measure}$ (Equation S8).

$$\text{Blockade}\left(Dose,t\right)_{\text{measure}} = \frac{Dose \times 100}{Dose + ED_{50}} \times \exp\left(-k \times \left(t - t_{\text{measure}}\right)\right) \qquad \text{S8}$$

In Equation S8, $t$ indicates the time in minutes after $t_{admin}$. By specifying $t = 0$ in Equation S8, we get the below function (Equation S9) for calculating the blockade produced by a given dose at $t_{admin}$ assuming no absorption phase.

$$\text{Blockade}\left(Dose\right)_{\text{admin}} = \frac{Dose \times 100}{Dose + ED_{50}} \times \exp\left(-k \times \left(0 - t_{\text{measure}}\right)\right) \qquad \text{S9}$$

By setting the volume of distribution $V$ to 1, we can substitute the dose input in the pkprofile function with the output of Equation S9 to enable estimation of time-blockade profiles.

##### 2.5. Calculating the absorption rate constant from time-to-peak MOR blockade

Equation S10 describes a general blockade-time profile of a bolus drug dose with an absorption phase and an elimination phase. The elimination rate constant is indicated by $k$, while the absorption rate constant is indicated by $k_a$.

$$\text{Blockade}\left(t\right) = \exp\left(-k \times t\right) - \exp\left(-k_a \times t\right) \qquad \text{S10}$$

Through derivation, we get the following expression for the rate of change in blockade at any given point in time ($t$):

$$\text{Blockade}'\left(t\right) = k_a \times \exp\left(-k_a \times t\right) - k \times \exp\left(-k \times t\right) \qquad \text{S11}$$

Because MOR blockade with a bolus dose of an opioid antagonist peaks at $t_{max}$, we know that

$$\text{Blockade}'\left(t_{\max}\right) = k_a \times \exp\left(-k_a \times t_{\max}\right) - k \times \exp\left(-k \times t_{\max}\right) = 0 \qquad \text{S12}$$

Rearranging the above equation, we get

$$k_a \times \exp\left(-k_a \times t_{\max}\right) = k \times \exp\left(-k \times t_{\max}\right) \qquad \text{S13}$$

This expression can be solved with the help of the Lambert W function ($W_{-1}$ branch):

$$-t_{\max} \times k_a \times \exp\left(-k_a \times t_{\max}\right) = -t_{\max} \times k \times \exp\left(-k \times t_{\max}\right) \qquad \text{S14}$$

$$W_{-1}\left(-t_{\max} \times k_a \times \exp\left(-k_a \times t_{\max}\right)\right) = W_{-1}\left(-t_{\max} \times k \times \exp\left(-k \times t_{\max}\right)\right) \qquad \text{S15}$$

$$-t_{\max} \times k_a = W_{-1}\left(-t_{\max} \times k \times \exp\left(-k \times t_{\max}\right)\right) \qquad \text{S16}$$

$$k_a = \frac{W_{-1}\left(-t_{\max} \times k \times \exp\left(-k \times t_{\max}\right)\right)}{-t_{\max}} \qquad \text{S17}$$

### 3. Results

#### 3.1. Naloxone

##### *3.1.1. Mu-opioid receptor blockade with intravenous naloxone*

Quantitative synthesis of data from PET [2,18] and dual-detector studies [16,19] of MOR blockade with intravenous naloxone was deemed appropriate due to the similarities in the protocols used across these studies. In both types of studies, a target dose of naloxone was first injected intravenously. Five minutes later, participants received an intravenous injection of radiolabeled [$^{11}$C]carfentanil, and positron emission from the brain was recorded for 60 minutes. The signal in the thalamus was used to calculate MOR blockade since this brain region has a very high density of mu-opioid receptors [20]. Blockade estimates in both types of studies were corrected for non-specific binding (i.e., non-receptor binding). The PET studies used the radiotracer signal in the occipital lobe as a reference for non-specific binding when correcting the blockade estimates [18] due to the small number of MOR in this region [20]. In contrast, the dual-detector studies used the thalamic radiotracer signal under 1.0 mg/kg intravenous naloxone as reference for non-specific binding because PET studies suggest that this dose is sufficient to block nearly all MOR in the thalamus [2,18].

*3.1.1.1. Difference in $ED_{50}$ between PET and dual-detector studies*

Modeling the dose-blockade relationship separately for PET ($RMSE$ = 7.88; $Pseudo\text{-}R^2$ = 0.92; Shapiro-Wilk test: $W$ = 0.85, $p$ = 0.16; Levene's test: $F_{1,4}$ = 4.54, $p$ = 0.10) and dual-detector studies ($RMSE$ = 7.38; $Pseudo\text{-}R^2$ = 0.96; Shapiro-Wilk test: $W$ = 0.91, $p$ = 0.28; Levene's test: $F_{1,8}$ = 0.08, $p$ = 0.78), we obtained $ED_{50}$ (SE) = 0.0050 mg/kg (0.0016) from PET data and $ED_{50}$ ($SE$) = 0.0018 mg/kg (0.0003) from dual-detector data. The difference in $ED_{50}$ between PET and dual-detector studies was not statistically significant ($\Delta ED_{50}$ = 0.0032 mg/kg, $SE$ = 0.0016, $t_{14}$ = 2.05, $p$ = 0.06; $RMSE$ = 7.57; $Pseudo\text{-}R^2$ = 0.95; Shapiro-Wilk test: $W$ = 0.91, $p$ = 0.10; Levene's test: $F_{1,14}$ = 1.69, $p$ = 0.21).

*3.1.1.2. Calculating the elimination rate constant*

Specifying an average blockade half-life of 110 minutes [16,21] in Equation S7, we get the following elimination rate constant:

$$k = \frac{\ln(2)}{110} = 0.006 \qquad \text{S18}$$

*3.1.1.3. Decrease in blockade between 45 and 65 minutes after naloxone administration*

Studies of MOR blockade with intravenous naloxone typically use the signal recorded between 45-65 minutes after naloxone administration to calculate blockade. Time series data from Kim et al. [16] indicate that [$^{11}$C]carfentanil activity and receptor blockade decreases linearly in this time frame, although blockade fluctuates considerably (Figure S1).

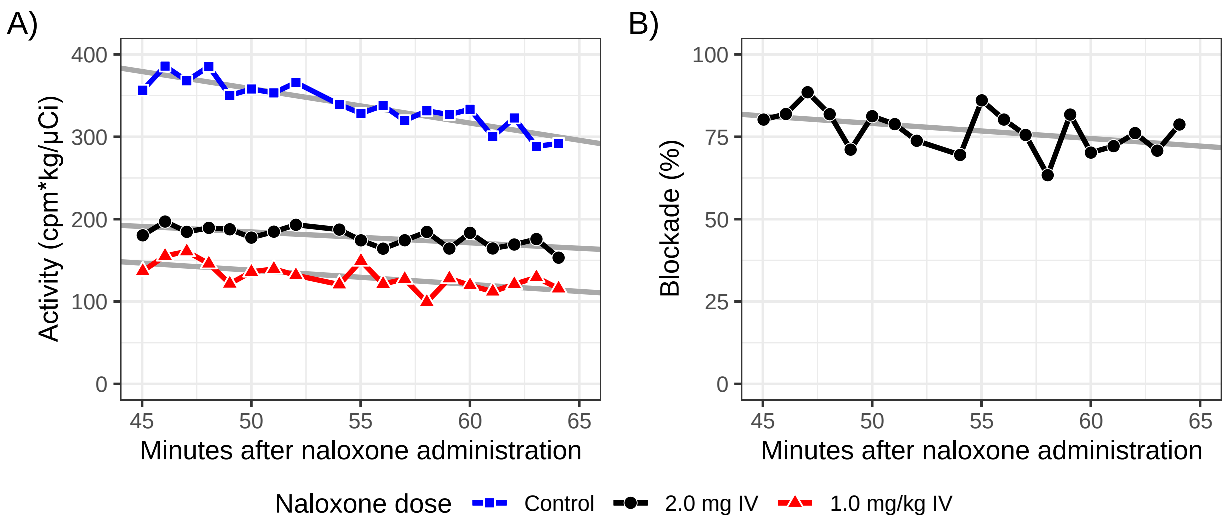


*Figure S1*. Decrease in [$^{11}$C]carfentanil activity (A) and blockade of [$^{11}$C]carfentanil binding (B) from 45-65 minutes after administration of intravenous (IV) naloxone. Blockade calculated according to Kim et al. [16]. Based on Kim et al. [16].

###### *3.1.1.4. Calculating the absorption rate constant*

By entering $t_{max}$ = 25 and $k$ = 0.006 in Equation S17, we can calculate the absorption rate constant $k_a$ that results in the blockade-time-profile peaking at 25 minutes after administration of intravenous naloxone (Equation S19).

$$k_a = \frac{W_{-1}\left(-25 \times 0.006 \times \exp\left(-0.006 \times 25\right)\right)}{-25} = 0.126 \qquad \text{S19}$$

#### 3.2. Other opioid antagonists

##### *3.2.1. Nalmefene*

Few studies have investigated receptor blockade with nalmefene. Figure S2 presents blockade estimates from a dual-detector study with a single dose of 1 mg and 0.001 mg/kg administered intravenously [16] and a PET study with 20 mg nalmefene administered orally as a single dose and daily over 7 days [22]. Based on data from the latter study, Kyhl et al. [23] estimate that a single dose of 20 mg oral nalmefene can maintain 60-90% blockade for up to 22-24 hours, with the blockade peaking at ~95% after 1-2 hours. Data on DOR and KOR occupancy with nalmefene is lacking. However, while nalmefene has low affinity to DOR, its high affinity to KOR indicates that it may not be possible to achieve full MOR blockade without producing substantial KOR occupancy (Table S1).

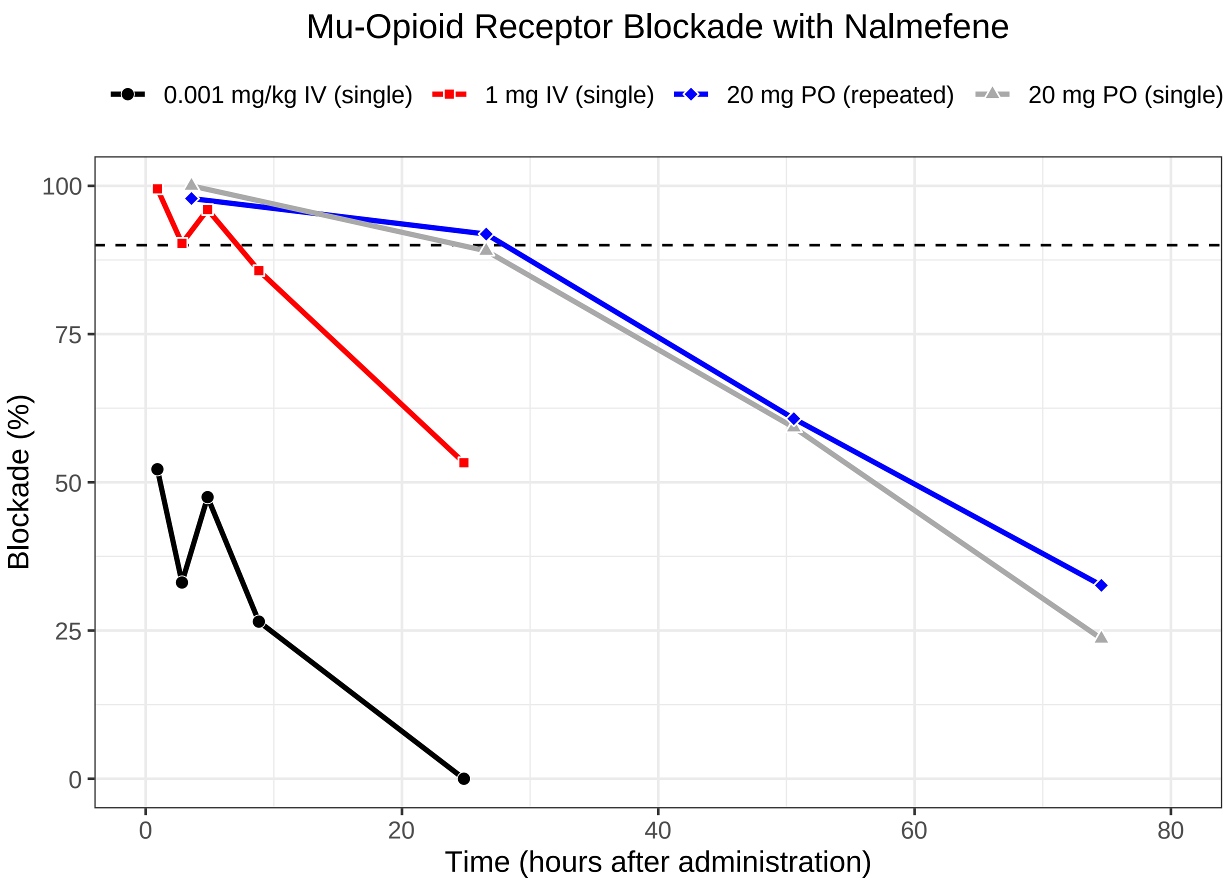


*Figure S2*. Mu-opioid receptor blockade with intravenous (IV) and oral (PO) nalmefene. Based on Kim et al. [16] and Ingman et al. [22]. The dashed horizontal line indicates full (90%) receptor blockade.

##### *3.2.2. GSK1521498*

GSK1521498 is an opioid antagonist not currently marketed for use in humans. The dose-blockade relationship for oral GSK1521498 is reported to be highly similar to that of oral naltrexone. When measured within 8 hours of administration, 0.4, 0.8, 1.5, 2, 6, 8, 10 and 50 mg oral produced 17, 38, 46, 59, 75, 85, 86 and 96% receptor blockade respectively [11]. Log-logistic modeling of these data (*RMSE* = 5.21; $Pseudo\text{-}R^2$ = 0.96; Shapiro-Wilk test: $W$ = 0.98, $p$ = 0.96; Levene's test: $F_{1,\,12}$ = 0.46, $p$ = 0.51) resulted in $ED_{50}$ (*SE*) = 1.51 mg (0.12) for MOR blockade with this antagonist. Our simulations suggests that oral GSK1521498 could achieve full MOR blockade while producing lower blockade of KOR than a similar dose of oral naltrexone due to its greater MOR-selectivity (Figure S3, Table S1). However, more data is needed to determine the onset and duration of full MOR blockade with this antagonist.

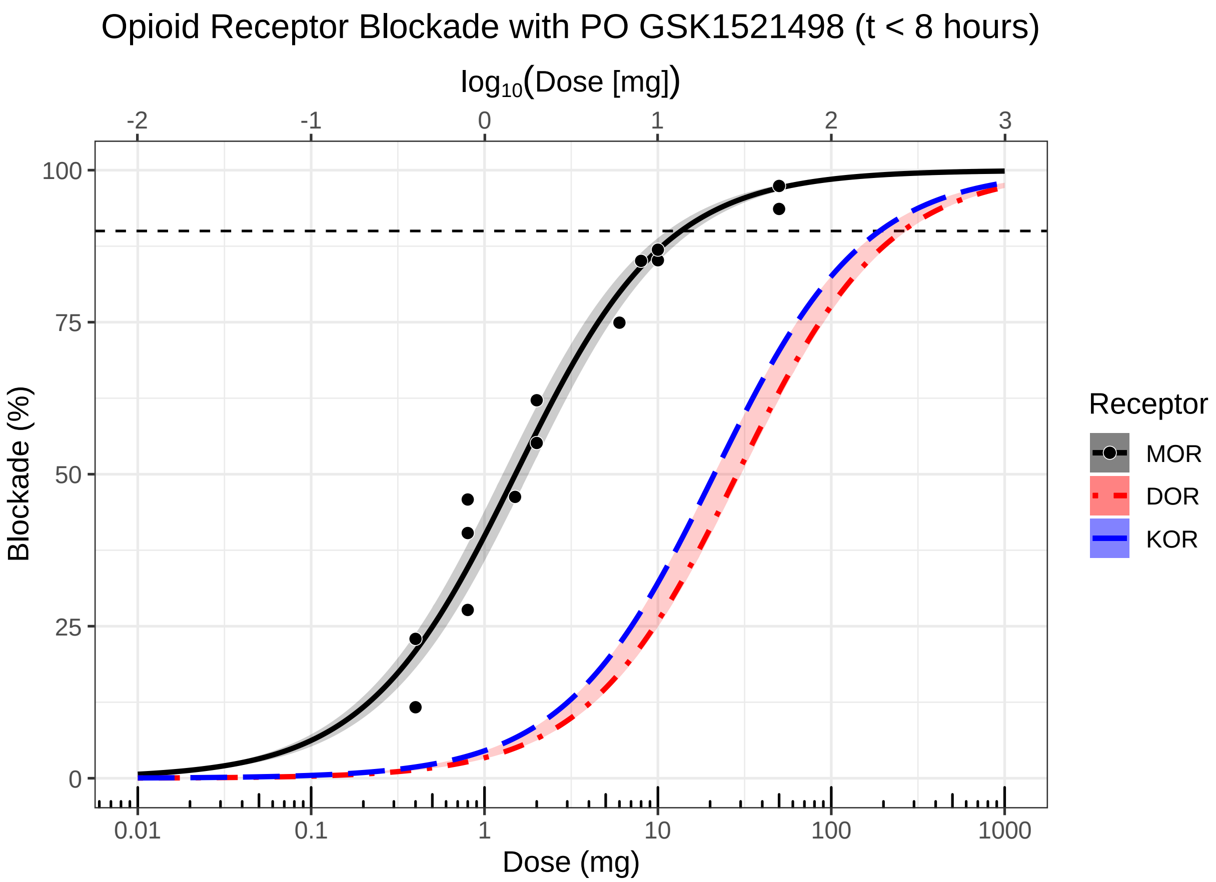


*Figure S3.* The effect of oral (PO) GSK1521498 dose on opioid receptor blockade within 8 hours of administration. The top x-axis displays $\log_{10}$-transformed doses while the bottom x-axis shows the corresponding untransformed doses. The dashed horizontal line indicates full (90%) receptor blockade. MOR blockade (solid black curve) is based on data (black dots) from Rabiner et al. [11]. DOR (dot-dashed red curve) and KOR blockade (long-dashed blue curve) were approximated from MOR blockade using the relative receptor affinities of GSK1521498 (Table S2). Semitransparent bands indicate 95% confidence band (black band), or range based on highest and lowest reported affinity ratio (blue and red bands; Table S1). The estimated $ED_{50}$ for MOR, DOR and KOR blockade was 1.51 ($SE$ = 0.12; see also [11]), 28.68 and 21.13 mg, respectively.

#### 4. Discussion

Reported MOR blockade estimates for large doses of naloxone and naltrexone are likely accurate due to ceiling effects [24]. However, estimates for lower doses and estimates from late in the elimination phase may be less precise due to large individual variability in receptor binding (e.g., [24]). Despite limited data, modeling of the dose-time-blockade relationship for intravenous naloxone and dose-blockade relationship for oral naltrexone was possible. The accuracy of our models can be validated against and improved with data from future PET studies.

Available time series data show a maximum reduction in [$^{11}$C]carfentanil activity from the control condition (i.e., no naloxone) on average 24.74 (SE = 1.24) minutes after administration of naloxone [1,16,17]. This gives an indication of the time it takes for naloxone to occupy a maximum amount of central MOR when administered as an intravenous bolus and can serve as a crude estimate of the time-to-peak ($t_{max}$) blockade. However, since none of these studies used continuous infusion of [$^{11}$C]carfentanil to maintain stable activity, it is also possible that the peak reduction in activity reflects a ceiling effect and that the time-to-peak naloxone concentration (and thus time-to-peak blockade) in the brain is longer than ~25 minutes. Continuous infusion of [$^{11}$C]carfentanil has been used in contemporary studies of MOR availability (for references, see [25]) and could potentially be used to obtain more accurate estimates of the time it takes for central MOR blockade to peak when naloxone is administered intravenously [26]. Because the model we present here is based on the currently available data, it does not account for variability in MOR blockade that may result from differences in [$^{11}$C]carfentanil administration method (i.e., bolus versus continuous infusion). However, our web application for naloxone enables users to alter the time-to-peak value used in the model.

The long-lasting MOR blockade produced by naltrexone is thought to result in part from metabolites of naltrexone that also act as antagonists [27]. However, some animal and human data suggest that naltrexone's main active metabolite 6β-Naltrexol has limited ability to pass the blood-brain barrier and block central agonist effects [28]. Considering naltrexone's long plasma terminal half-life (96 hours [29]), it is possible that the prolonged blockade is facilitated by non-specific binding retention and/or slow release of naltrexone from central receptors. Regardless of cause, naltrexone's long central blockade half-life implies that long intersession intervals are necessary to avoid carry-over effects from residual blockade.

We have primarily discussed the usefulness of intravenous naloxone and oral naltrexone in basic human research on the brain's opioid system. Intranasal formulations of naloxone are also marketed for human use. These are noninvasive and easy to apply, and may therefore represent a viable alternative to intravenous naloxone and oral naltrexone. In a PET study by

Johansson et al. [21], 4 mg of intranasal naloxone produced 85% MOR blockade and maintained this level of blockade for at least 20 minutes. However, there was large individual variability in peak MOR blockade, perhaps due to differences in nasal physiology and administration technique [30]. The observed time-to-peak MOR blockade was ~40 minutes. Due to the slow absorption phase and risk of suboptimal administration, higher intranasal doses than 4 mg may be necessary to achieve full blockade more quickly and across all subjects. We therefore recommend using intravenous naloxone and oral naltrexone over intranasal naloxone to block the endogenous opioid system in basic human research. Compared to intranasal administration, intravenous administration requires smaller doses of naloxone to quickly produce high MOR blockade and is therefore less costly. Oral naltrexone likely results in less variable MOR blockade than intranasal naloxone due to the greater reliability of the oral administration method.

The introduction of opioid antagonist drugs like naloxone and naltrexone has enabled researchers to investigate the causal role of the endogenous opioid system in psychological processes and behavior in humans. In studies with human participants, researchers are also probing the mu-opioid system through naturally occurring variation in the mu-opioid receptor gene [31] and functional imaging of mu-opioid receptor availability with PET [32]. These methods are however very costly due to the need for large samples (e.g., candidate gene studies) or specialized radiolabeling setups (e.g., PET). Psychopharmacological studies with opioid antagonists can also become expensive due to suboptimal dosing schedules. Our web applications for intravenous naloxone (https://martintrostheim.shinyapps.io/planoxone/) and oral naltrexone (https://martintrostheim.shinyapps.io/plantrexone/) enable researchers to easily select a dosing schedule that minimizes drug cost while still achieving the desired level of blockade.

#### References

1. Bice AN, Wagner HN, Frost JJ, Natarajan TK, Lee MC, Wong DF, et al. Simplified Detection System for Neuroreceptor Studies in the Human Brain. J Nucl Med. 1986;27:184–191.
2. Frost JJ, Wagner HNJr, Dannals RF, Ravert HT, Links JM, Wilson AA, et al. Imaging Opiate Receptors in the Human Brain by Positron Tomography. Journal of Computer Assisted Tomography. 1985;9:231–236.
3. Zhang Y, Fox GB. PET imaging for receptor occupancy: meditations on calculation and simplification. Journal of Biomedical Research. 2012;26:69–76.

4. Kelly E, Mundell SJ, Sava A, Roth AL, Felici A, Maltby K, et al. The opioid receptor pharmacology of GSK1521498 compared to other ligands with differential effects on compulsive reward-related behaviours. Psychopharmacology. 2015;232:305–314.
5. Peng X, Knapp BI, Bidlack JM, Neumeyer JL. Pharmacological Properties of Bivalent Ligands Containing Butorphan Linked to Nalbuphine, Naltrexone, and Naloxone at μ, δ, and κ Opioid Receptors. J Med Chem. 2007;50:2254–2258.
6. Toll L, Berzetei-Gurske I, Polgar W, Brandt S, Adapa I, Rodriguez L, et al. Standard Binding and Functional Assays Related to Medications Development Division Testing for Potential Cocaine and Opiate Narcotic Treatment Medications. NIDA Research Monograph. 1998;178:440–466.
7. Konoura K, Fujii H, Imaide S, Gouda H, Hirayama S, Hirono S, et al. Transformation of naltrexone into mesembrane and investigation of the binding properties of its intermediate derivatives to opioid receptors. Bioorganic & Medicinal Chemistry. 2015;23:439–448.
8. Schüllner F, Meditz R, Krassnig R, Morandell G, Kalinin VN, Sandler E, et al. Synthesis and Biological Evaluation of 14-Alkoxymorphinans. Part 19. Helvetica Chimica Acta. 2003;86:2335–2341.
9. Wentland MP, Lu Q, Lou R, Bu Y, Knapp BI, Bidlack JM. Synthesis and opioid receptor binding properties of a highly potent 4-hydroxy analogue of naltrexone. Bioorganic & Medicinal Chemistry Letters. 2005;15:2107–2110.
10. Bart G, Schluger JH, Borg L, Ho A, Bidlack JM, Kreek MJ. Nalmefene Induced Elevation in Serum Prolactin in Normal Human Volunteers: Partial Kappa Opioid Agonist Activity? Neuropsychopharmacology. 2005;30:2254–2262.
11. Rabiner EA, Beaver J, Makwana A, Searle G, Long C, Nathan PJ, et al. Pharmacological differentiation of opioid receptor antagonists by molecular and functional imaging of target occupancy and food reward-related brain activation in humans. Molecular Psychiatry. 2011;16:826–835.
12. Rohatgi A. WebPlotDigitizer. 2021.
13. Burda BU, O'Connor EA, Webber EM, Redmond N, Perdue LA. Estimating data from figures with a Web-based program: Considerations for a systematic review. Research Synthesis Methods. 2017;8:258–262.
14. Drevon D, Fursa SR, Malcolm AL. Intercoder Reliability and Validity of WebPlotDigitizer in Extracting Graphed Data. Behav Modif. 2017;41:323–339.
15. Moeyaert M, Maggin D, Verkuilen J. Reliability, Validity, and Usability of Data Extraction Programs for Single-Case Research Designs. Behav Modif. 2016;40:874–900.
16. Kim S, Wagner HN, Villemagne VL, Kao P-F, Dannals RF, Ravert HT, et al. Longer Occupancy of Opioid Receptors by Nalmefene Compared to Naloxone as Measured In Vivo by a Dual-Detector System. J Nucl Med. 1997;38:1726–1731.
17. Lee MC, Wagner HN, Tanada S, Frost JJ, Bice AN, Dannals RF. Duration of Occupancy of Opiate Receptors by Naltrexone. J Nucl Med. 1988;29:1207–1211.
18. Mayberg HS, Frost JJ. Opiate Receptors. In: Frost JJ, Wagner Jr HN, editors. Quantitative Imaging: Neuroreceptors, Neurotransmitters, and Enzymes, New York: Raven Press, Ltd.; 1990.
19. Villemagne VL, Frost JJ, Dannals RF, Lever JR, Tanada S, Natarajan TK, et al. Comparison of [11C]diprenorphine and [11C]carfentanil in vivo binding to opiate receptors in man using a dual detector system. European Journal of Pharmacology. 1994;257:195–197.

20. Hirvonen J, Aalto S, Hagelberg N, Maksimow A, Ingman K, Oikonen V, et al. Measurement of central μ-opioid receptor binding in vivo with PET and [11C]carfentanil: a test–retest study in healthy subjects. European Journal of Nuclear Medicine and Molecular Imaging. 2008;36:275.
21. Johansson J, Hirvonen J, Lovró Z, Ekblad L, Kaasinen V, Rajasilta O, et al. Intranasal naloxone rapidly occupies brain mu-opioid receptors in human subjects. Neuropsychopharmacology. 2019;44:1667–1673.
22. Ingman K, Hagelberg N, Aalto S, Någren K, Juhakoski A, Karhuvaara S, et al. Prolonged Central μ-Opioid Receptor Occupancy after Single and Repeated Nalmefene Dosing. Neuropsychopharmacology. 2005;30:2245–2253.
23. Kyhl L-EB, Li S, Faerch KU, Soegaard B, Larsen F, Areberg J. Population pharmacokinetics of nalmefene in healthy subjects and its relation to μ-opioid receptor occupancy. British Journal of Clinical Pharmacology. 2016;81:290–300.
24. Weerts EM, Kim YK, Wand GS, Dannals RF, Lee JS, Frost JJ, et al. Differences in δ- and μ-Opioid Receptor Blockade Measured by Positron Emission Tomography in Naltrexone-Treated Recently Abstinent Alcohol-Dependent Subjects. Neuropsychopharmacology. 2008;33:653–665.
25. Blecha JE, Henderson BD, Hockley BG, VanBrocklin HF, Zubieta J-K, DaSilva AF, et al. An updated synthesis of [11C]carfentanil for positron emission tomography (PET) imaging of the μ-opioid receptor. Journal of Labelled Compounds and Radiopharmaceuticals. 2017;60:375–380.
26. Kao P-F, Kim S, Wagner HN, Lever JR, Ravert HT, Dannals RF. Assessing neuroreceptor occupancy by continuous infusion of carbon-11 labeled radioligands. European Journal of Nuclear Medicine. 1996;23:141–144.
27. Schmidhammer H. Opioid Receptor Antagonists. In: Ellis GP, Luscombe DK, Oxford AW, editors. Progress in Medicinal Chemistry, vol. 35, Elsevier; 1998. p. 83–132.
28. Yancey-Wrona J, Dallaire B, Bilsky E, Bath B, Burkart J, Webster L, et al. 6β-Naltrexol, a Peripherally Selective Opioid Antagonist that Inhibits Morphine-Induced Slowing of Gastrointestinal Transit: An Exploratory Study. Pain Medicine. 2011;12:1727–1737.
29. Verebey K, Volavka J, Mule SJ, Resnick RB. Naltrexone: Disposition, metabolism, and effects after acute and chronic dosing. Clinical Pharmacology & Therapeutics. 1976;20:315–328.
30. Pires A, Fortuna A, Alves G, Falcão A. Intranasal Drug Delivery: How, Why and What for? J Pharm Pharm Sci. 2009;12:288–311.
31. Mague SD, Blendy JA. OPRM1 SNP (A118G): Involvement in disease development, treatment response, and animal models. Drug and Alcohol Dependence. 2010;108:172–182.
32. Henriksen G, Willoch F. Imaging of opioid receptors in the central nervous system. Brain. 2008;131:1171–1196.
